# Accurate *de novo* transcription unit annotation from run-on and sequencing data

**DOI:** 10.1101/2025.02.12.637853

**Authors:** Paul R. Munn, Jay Chia, Charles G. Danko

## Abstract

Functional element annotations are critical tools used to provide insight into the molecular processes governing cell development, differentiation, and disease. Run-on and sequencing assays measure the production of nascent RNAs and can provide an effective data source for discovering functional elements. However, the accurate inference of functional elements from run-on sequencing data remains an open problem because the signal is noisy and challenging to model. Here we investigated computational approaches that convert run-on and sequencing data into annotations representing transcription units, including genes and non-coding RNAs. We developed a convolutional neural network, called <u>c</u>onvolutional discovery of <u>g</u>ene <u>a</u>natomy using <u>P</u>RO-seq (CGAP), trained to identify different anatomical features of a transcription unit, which were then stitched together into transcript annotations using a hidden Markov model (HMM). Comparison with existing methods showed a significant performance improvement using our novel CGAP-HMM approach. We developed a voting system that ensembles the top three annotation strategies, resulting in large and significant improvements in transcription unit annotation accuracy over the best performing individual method. Finally, we also report a conditional generative adversarial network (cGAN) as a generative approach to transcription unit annotation that shows promise for further development. Collectively our work provides novel tools for *de novo* transcription unit annotation from run-on and sequencing data that are accurate enough to be useful in many applications.

## 1. INTRODUCTION

Genomes encode a diverse collection of functional elements that provide the instruction manual for cell development, differentiation, and homeostasis (Hnisz *et al*., 2016; Lupiáñez *et al*., 2015). Annotating the location and conditional activity of functional elements that make up this instruction manual is a critical goal of modern genetics. Genome annotations are tools that are crucial for understanding, cloning, and mapping genome sequences. In humans and mice, protein-coding and non-coding mRNAs are annotated through painstaking efforts of large consortia, such as RefSeq (O’Leary *et al*., 2016) and GENCODE (Harrow *et al*., 2012) or built using population-scale RNA-seq data (Pertea *et al*., 2018). In addition to mRNA encoding genes, a wide variety of non-coding functional elements also serve important roles.

Non-coding functional elements in the human and mouse genomes have been annotated by the concerted efforts of the ENCODE and Epigenome Roadmap consortia, which use a combination of assays (e.g., ChIP-seq, DNase, mRNA-seq) to identify active genomic regions across numerous cell types and tissues (Ernst and Kellis, 2012; Hoffman *et al*., 2012). Efforts are underway to extend these functional annotations to common agricultural, veterinary, and plant species (Koepfli *et al*., 2015; Darwin Tree of Life Project Consortium, 2022). Genome annotation efforts are likely to intensify in coming years as moonshot projects designed to sequence the entire tree of life come to fruition (Lewin *et al*., 2018). Yet applying the approaches pioneered in human and mouse to genomes across the tree of life remains a formidable challenge due to the resources and expertise required.

An alternative approach to the integration of multiple functional assays is to focus on a single experiment which maximizes information about genome function in a sample. RNA polymerase II (Pol II), the RNA polymerase which transcribes all protein-coding genes and most non-coding RNAs, leaves characteristic patterns that can be used to distinguish a wide variety of functional elements (Wang *et al*., 2019, 2022). Most transcription units produced by Pol II are rapidly degraded by the nuclear exosome complex (Ntini *et al*., 2013; Andersson *et al*., 2015), and therefore identifying them requires genome-wide measurements that directly capture RNA polymerase, such as precision nuclear run-on and sequencing (PRO-seq) or related assays (Kwak *et al*., 2013). PRO-seq data, when viewed in a genome browser, has characteristic shapes that are informative of the molecular function of a particular DNA sequence. Indeed, existing approaches have been proposed using a variety of machine learning and probabilistic modelling methods to interpret distinctive patterns of Pol II with substantial success (Azofeifa and Dowell, 2017; Wang *et al*., 2019).

Existing computational approaches to interpret PRO-seq data have largely focused on the discovery of either transcription initiation regions (TIRs) or the boundaries of active transcription units. Here we report a machine learning strategy which identifies patterns of RNA polymerase associated with several anatomical landmarks in an active transcription unit using a multi-task convolutional neural network. By combining the output of these shapes with hidden Markov models (HMMs), we developed a general strategy that discovers the boundaries of a wide variety of transcription units. Additionally, we developed an ensemble classifier using our new CNN-HMM-based method and two existing tools, groHMM (Chae *et al*., 2015) and T-units (Danko *et al*., 2018), in such a way as to overcome weaknesses of each approach. Lastly, we explored a strategy that uses a conditional generative adversarial network (cGAN) to directly map PRO-seq signal to corresponding annotations. These analysis tools confirm the availability of additional shapes in PRO-seq signal that we leverage here to improve the annotation of genomes using a single experimental assay.

## 2. METHODS

### 2.1 Label acquisition and processing

Training labels for the CNN and cGAN were obtained from the same datasets and then processed in different ways. We begin by outlining the initial processing steps that are common to the two methods and then discuss the differences separately.

K562 PRO-seq and GRO-seq datasets G3, G5, G6, and G7 were used for training, with GRO-seq dataset G2 used for validation, and PRO-seq dataset G1 used as a holdout dataset (see Table 1). Possible overfitting to lab specific technical variation was reduced by selecting each dataset from separate labs. Analysis done on the same datasets in (Wang *et al*., 2019) showed that RPKM normalized read counts were highly correlated.

**Table 1:** Data sources.

| Data ID | Cell line | GEO | Organism | Protocol | Train/<br>Test | Reference |
| --- | --- | --- | --- | --- | --- | --- |
| G1 | K562 | GSM1480327 | Human | PRO-seq | Test | (Core *et al.*, 2014) |
| G2 | K562 | GSM1480325 | Human | GRO-seq | Test | (Core *et al.*, 2014) |
| G3 | K562 | GSM3452725 | Human | PRO-seq | Train | (Wang *et al.*, 2019) |
| G5 | K562 | GSM2361442<br>GSM2361443 | Human | PRO-seq | Train | (Vihervaara *et al.*, 2017) |
| G6 | K562 | GSM2545324 | Human | PRO-seq | Train | (Dukler *et al.*, 2017) |
| G7 | K562 | GSM2545325 | Human | PRO-seq | Train | (Dukler *et al.*, 2017) |
| GM | GM12878 | GSM1480326 | Human | GRO-seq | Test | (Core *et al.*, 2014) |
| GH | HCT116 | GSM1304424<br>GSM1304425 | Human | GRO-seq | Test | (Allen *et al.*, 2014) |
| CD4 | CD4 | GSM2265095 | Human | PRO-seq | Test | (Danko *et al.*, 2018) |
| MCF7 | MCF-7 | GSM1014637<br>GSM1014638<br>GSM1014637<br>GSM1014638<br>GSM1014639 | Human | GRO-seq | Test | (Danko *et al.*, 2013;<br>Hah *et al.*, 2011) |
| H1 | HELA | GSE62046 | Human | GRO-seq | Test | (Andersson *et al.*, 2014) |
| K562 GRO-cap | K562 | GSM1480321 | Human | GRO-cap | Train | (Core *et al.*, 2014) |
| GM GRO-cap | GM12878 | GSM1480323 | Human | GRO-cap | Test | (Core *et al.*, 2014) |
| DNase UW | K562 | NA | Human | DNase-seq | Train | (ENCODE Project Consortium, 2012) |
| DNase Duke | K562 | NA | Human | DNase-seq | Train | (ENCODE Project Consortium, 2012) |
| Poly-A | K562 | NA | Human | RNA-seq | Train | (ENCODE Project Consortium, 2012) |
| GENCODE | K562 | NA | Human | Multiple | Train | (Frankish *et al.*, 2019) |

We began by selecting different regions that could serve as informative training loci. Specifically, we chose regions with high PRO-seq signal, defined by previous studies as windows with more than 3 PRO-seq reads within 100 bp on a single strand or at least one read within 1000 bp on both positive and negative strands (Wang *et al*., 2019). These criteria exclude regions such as the centromeric region that show little to no PRO-seq signal and thus provide little value as training labels. We expanded each of these informative regions by 20 Kb at either end into the low signal regions, so that the negative training labels (i.e. regions labeled as non-transcribed) would partially include low coverage PRO-seq signal.

To choose training labels from these informative positions we began by constructing a set of high confidence GRO-cap sites – these are GRO-cap sites that intersect with both DNase UW, and DNase Duke peak calls provided by ENCODE (see data access). These high-confidence GRO-cap sites were further divided into stable and unstable groups, using definitions provided in previous publications (Tome *et al*., 2018).

To define gene bodies, we used GENCODE annotations that overlap the region between high confidence GRO-cap sites and polyadenylation sites, and that also fall within an informative region. This subset of high confidence GENCODE annotations constitutes the gene body labels.

To build a set of transcription unit start labels we took the intersection of the high confidence GRO-cap sites with a 1 Kb window centered on the 5’ end of each of the gene body labels from the previous step. Likewise, for the set of transcription end labels we took the intersection of the polyadenylation sites (see Table 1) with a 1 Kb window centered on the 3’ end of each of the gene body labels. The set of labels referencing after transcription units was built simply by extending a 10 Kb window downstream of each of the end labels. To construct a set of labels for non-transcribed regions we subtracted all the labels for gene bodies, start sites, end sites, and after transcription unit regions from the genome reference. Each of the above sets of labels was subsequently divided into two sets – one for the plus strand and one for the minus strand. We used the set of high confidence GRO-cap sites as the set of TSS labels but kept this as non-strand specific.

To further filter the sets of labels, we removed from the informative positions defined at the outset any place where we have an active GENCODE annotation and no T-units prediction, and any place where we have a T-units prediction, but no active GENCODE annotation. The rational for this is T-units attempts to predict positions that are transcribed, so loci with a T-unit call (but no active GENCODE annotation) are enriched for unannotated lincRNAs and would likely confuse discriminative model training.

Finally, we compiled a list of places where each of the transcription unit bodies, transcription unit starts, transcription unit ends, after transcription unit positions, TSSs, and non-transcribed regions intersect with the filtered informative positions dataset. We associated the appropriate label with each of these intersecting regions and calculated the coverage for that region. When training, we select from these labels at random, assigning higher probability of selection to those regions with higher coverage.

When selecting the proportions of training labels to train on, we opted not to keep the exact distribution of labels within the genome. This is because the non-transcribed labels dominate the genome. We increased the representation of training labels from transcribed regions. The following proportions were used: non-transcribed training samples: 40%; transcription unit body samples: 30%; transcription unit start site samples: 5%; transcription unit end site samples: 5%; after transcription unit region samples: 5%; TSS samples: 5%; transcription unit stable start site samples: 5%; transcription unit unstable start site samples: 5%.

### 2.2 cGAN training labels

Training samples for the GAN were approached somewhat differently. Rather than giving an entire region a single label (i.e. the label associated with the data at the center of the region) as we did above, we divided the training region into 50bp bins and assigned labels of gene body, after-gene region, and transcription start sites to each bin in the training region. Labels were assigned based on both GENCODE annotations and consensus annotations (as defined in section 2.7 *Building a consensus for CGAP-HMM, groHMM, and T-units* below). Consensus annotations performed better and are reported in section 2.8 *Genome-wide accuracy metrics for transcription unit identification* below.

By way of example, the distinction between the labeling approaches can be thought of in terms of image processing of a street scene. For the CNN, the entire scene might be given a label such as “red car,” which is found at the center of the image, whereas for the GAN the training is treated as more of an image segmentation task where each object in the scene (cars, people, street signs, road and sidewalk, buildings, etc.) is given a label along with its location. In other words, labels are generated in a similar fashion to those for the CNN but presented to the GAN as a vector of values.

### 2.3 Transcription unit inference using a CNN and an HMM

We approached the problem of transcription unit inference as a two-step process. First, we predicted the locations of features such as transcription starts, transcription ends, gene body transcription, and after gene transcription based on patterns evident in the PRO-seq signal. It should be noted here that these features are subject to much of the same transcriptional noise as the original PRO-seq signal, and so cannot be used directly to infer transcriptional elements. Instead, we employ a second step where we use these features as emissions in a hidden Markov model, effectively smoothing over random spikes in the signal and/or short regions of low or zero signal. After smoothing, we infer the beginning and end of transcriptional elements as continuous regions of chromatin with well-defined start and end points.

### 2.4 CNN architecture and implementation

Architecture: The CNN consists of an initial input layer with dimensions of 1024 by 2. Prior to using the PRO-seq data as input, we bin the reads into 50 bp regions. Thus, the 1024 width of the input layer corresponds to 1024 x 50 bp, or 51.2 Kb. The height of 2 for the input corresponds to the signal on the plus and minus strands.

Following the input layer are five convolutional layers, each using a rectified linear unit (ReLU) activation function, and a 0.2x dropout. Each convolutional layer is separated by a batch normalization layer and MaxPool layer that subsamples the width dimension by a factor of 2. The final convolutional layer is flattened and passed to a fully connected layer of 256 ReLU nodes (also with 0.2x dropout), which then connects to 15 logistic regression nodes that output the probabilities for each of the 15 labels.

Training: Labeled regions were selected from a pool of approximately 740,000 labels in batches of 128. Although selection is at random, higher probability is given to regions of the PRO-seq signal with higher coverage. Once a training region has been selected, a random point within that region is chosen as the center of point of a 51.2 Kb window, that is then used as input to the CNN. It should be noted that in the case of a small label, such as a TSS of 140 bp in width, it is narrow enough that it will remain at the approximate center of the window. However, in the case of a much wider label, such as a 200 Kb gene body, the center point of the window could effectively be anywhere within the gene.

We experimented with hyper-parameter settings (convolution filter width, number of convolution layers, dropout rate, etc.) and tested these models on the validation set. We found that an input layer with 1024 nodes produced the best results, but found no other performance differences, so long as the network was sufficiently large.

The network was trained using the Adam optimizer, an adaptive learning rate optimization algorithm, specifically designed for training deep neural networks (Kingma and Ba, 2014). Training ran for 4,800 epochs which took approximately 5 days on an NVIDIA Tesla TITAN X (Pascal) GPU.

Evaluation: The “roc_curve” and “precision_recall_curve” functions from the scikit-learn metrics Python library (Pedregosa *et al*., 2011) were used to evaluate the performance of each version of the CNN. While these functions do not give an absolute measure of how well a particular CNN is able to predict transcription units (due to uncertainty in assigning training labels), they do provide a metric for comparing relative performance of one CNN against another and thus enable us to select the “best” one. For both functions, we passed two vectors as parameters: a vector indicating the training label for the region being evaluated, and a vector of the predicted probabilities produced by the CNN for each possible label. For the roc_curve function we plotted the fraction of true positives vs. the fraction of false positives, for each of the 15 labels, at various thresholds for the label probability. We then calculated the area under the receiver operating characteristic curve (AUC) as the performance metric (larger values indicate a higher proportion of true positives vs. false positives). For the precision_recall_curve function we plotted the precision of our classifier vs. its recall (high precision indicates a low false positive rate and high recall indicates a low false negative rate). Again, we calculated the AUC and use this as a metric to assess performance (larger values indicate larger numbers of accurate results).

### 2.5 Implementation of HMM

The primary goal of the hidden Markov model was to smooth CNN predictions into contiguous transcription unit predictions. RNA Pol II continues to actively elongate beyond the end of gene bodies resulting in the CNN predicting continued transcription in this after gene region. We took this into account by building a three-state model for the HMM: the first state is non-transcribed, which changes to the second state of transcribed with a probability learned by the model upon encountering an increase in the PRO-seq signal and / or a start site (also predicted by the CNN). From this second, transcribed state, the HMM transitioned to the third state of ‘after-gene transcription’, again using a probability learned by the model, before finally transitioning back to the non-transcribed state.

We explored alternative HMM structures. HMMs with only two states (non-transcribed and transcribed) performed well when finding the beginnings of transcription units but tended to overestimate the length of these annotations. Introducing a third state attempts to enable the model to compensate for the after-gene transcription and thus find a more accurate end point for each annotation. We also explored models that used more of the 15 different CNN labels. Using up to 11 of the 15 sets of predicted labels as emissions combined with a 5 state (representing non-transcribed – transcription unit start – transcription unit body – transcription unit end – after transcription unit) HMM, we experimented with various discrete and continuous emissions distributions, and the inclusion of different sets of covariates (start labels, end labels, and after transcription unit labels).

In some HMM architectures, covariates were used as a potential signal for a state change (for example, in the best performing model described in the text, start labels assigned by the CNN were used as transition probabilities to signal a change from non-transcribed to transcribed states). For each covariate used, the maximum value of the covariate within each 50 bp bin, clamped to the zero to one interval, was used as a transition probability between the appropriate states.

Ultimately, the relatively simple HMM, introduced above, using three states, the plus and minus transcription unit body labels as emissions, and plus and minus transcription unit start loci as covariates to influence the transition probability from non-transcribed to transcription unit body achieved the best performance for transcription unit annotation. All HMMs were implemented using the QHMM package (https://github.com/andrelmartins/QHMM) in R.

In all cases we used the Baum-Welch algorithm (Baum and Petrie, 1966) to perform expectation maximization. Evaluation for each HMM was performed using the metrics in section *2.8 (Genome-wide accuracy metrics for transcription unit identification)*.

### 2.6 Implementation of cGAN

The cGAN we used was based heavily on a TensorFlow 2.0 (Abadi *et al*., 2016) implementation of the pix2pix cGAN written by (Isola *et al*., 2017). The TensorFlow implementation, copywritten in 2019 by “The TensorFlow Authors” is licensed for use under the Apache License, Version 2.0 (https://www.apache.org/licenses/LICENSE-2.0)

Architectures: The cGAN consists of two neural networks - a generator and a discriminator. They are described here with the modifications necessary for building a model that operates on the plus and minus strands of a PRO-seq dataset.

The generator was built as a modified U-NET configuration (Ronneberger *et al*., 2015) – this is an encoder-decoder configuration with skip connections between the encoder layers and the corresponding decoder layers. The encoder is built from multiple convolution layers, each using a ReLU activation and each separated by a batch normalization layer. The decoder is built from multiple transposed convolution layers, also using a ReLU activation layer and each separated by a batch normalization layer, with dropout applied to the first three layers.

The discriminator is modeled on a PatchGAN (Li and Wand, 2016) where each “patch” of the final output layer classifies a larger region of the input layer. Other than this, it has a similar configuration to the encoder above, with multiple convolutional layers, each using a ReLU activation, and each separated by a batch normalization layer.

Training: for a typical GAN, training takes place by alternately running the generator for a single epoch and then the discriminator for a single epoch, and continuing in this fashion so as to prevent one network getting too far ahead of the other (Goodfellow *et al*., 2014).

Each example input consisted of a two-row matrix containing the PRO-seq signal from a region approximately 1.6 Mb wide (row one of the matrix was the plus strand and row two was the minus strand). For each region the generator generates an output. The discriminator is given this generated sample along with the corresponding PRO-seq signal as its first input. For the discriminator’s second input we use the input PRO-seq signal and the target annotation for same region the PRO-seq was taken from.

We can now calculate the losses for the generator and the discriminator. The loss for the discriminator is the accuracy with which it can distinguish the generated sample from the target annotation. The loss for the generator is based on its ability to generate a sample that the discriminator classifies as a real (target) annotation (i.e. how well it can fool the discriminator).

Training ran for approximately 1,000 epochs which took approximately 2 days on an NVIDIA Tesla TITAN X (Pascal) GPU.

Evaluation: Assessing the performance of a GAN by looking at the losses for the generator and discriminator can be misleading, since they fluctuate as training alternates between improving one and then the other network. As mentioned above, one problem with GAN training is one network getting too far ahead of the other: a symptom of this is the loss for one network gets very low – this happens when one network is dominating the other and is an indicator that the training of the combined network has run into problems. The authors of the Isola paper created an L1 loss for the generator that should go down as training progresses, but ultimately the measure of the combined networks performance are the metrics shown in section *2.8 (Genome-wide accuracy metrics for transcription unit identification)*.

### 2.7 Building a consensus for CGAP-HMM, groHMM, and T-units

To combine the strengths of various methods for transcription unit identification, we developed an ensemble approach to construct consensus transcription unit annotations using predictions from CGAP-HMM, groHMM, and T-units. Our approach works as follows:

- Remove small fragments (<101 bp) from each annotation dataset.
- Find the intersection of all annotations in each pair of datasets.
- Combine these intersections into a single dataset.
- Add back all transcription units from any method which does not intersect a transcription unit in the combined set. This will enable us to keep predictions made by only one method.

Examination of the results in a genome browser showed obvious cases where an annotation had been predicted by one method, but only partially predicted by the other two. However, these cases were excluded from the results because of the very small intersection of the partial predictions. To overcome this, we modified the final step of the process above to add back transcription units from any method that does not intersect a transcription unit in the combined set by more than 10%.

### 2.8 Genome-wide accuracy metrics for transcription unit identification

Two of the metrics used in the evaluation of transcription unit boundaries require special explanation. The first of these measures the accuracy with which each method recovered the boundaries of gene annotations in gene-dense regions. Two types of error can arise in this measurement: either TU identification methods can erroneously break annotated genes into smaller predicted gene fragments, or alternatively a single transcription unit prediction can erroneously merge two or more separate gene annotations (**Supplemental figure S1A**). While the optimal method would get both values as low as possible, it should be noted that incorrect parameter tuning can result in one value decreasing at the expense of the other. For example, it would be easy to decrease the count of annotations disassociated by increasing the number of annotations merged (or vice versa), but this should be avoided.

We also used the Transcription Unit Accuracy (TUA) metric defined by (Chae *et al*., 2015) to evaluate TU annotation accuracy. The TUA metric implemented in the groHMM software package (Chae *et al*., 2015) quantifies TU annotation error in either over-or under-estimating the length of gene annotations. In brief, the TUA metric measures the accuracy of the 5’ and 3’ end of transcription unit predictions by dividing each prediction into three sections (upstream of the TSS, within the transcribed region, and downstream of the polyadenylation cleavage site). It then does the same for the ground truth annotations (we defined a set of GENCODE annotated genes which have a GRO-cap peak near the transcription start site and a poly-A RNA-seq peak near the polyadenylation cleavage site). The TUA method will penalize predicted annotations that begin before the TSS of the ground truth annotation, or that end before the polyadenylation cleavage site (**Supplemental figure S1B**). Since Pol II continues to transcribe past the polyadenylation cleavage site (Kwak *et al*., 2013), TUA does not penalize for extending annotations past the polyadenylation cleavage site. The TUA metric ranges between 0 and 1, and values near 1 indicate higher agreement with the ground truth.

## 3. RESULTS

### 3.1 A multi-class CNN can identify distinct anatomical landmarks in a transcription unit

Our first goal was to develop a strategy to discover anatomical landmarks in active genes. Motivated by the success of dREG and related tools (Wang *et al*., 2019), we focused on using a supervised machine learning approach to interpret patterns in PRO-seq data at multiple scales. We developed a multi-class convolutional neural network (CNN) trained to recognize the pattern of RNA polymerase associated with fifteen distinct labels, which represent different parts of the anatomy of a transcription unit. Labels include broad transcription initiation regions (TIRs) which initiate new Pol II, the gene body of active transcription units, the start position of stable transcription units, the start of exosome sensitive unstable transcription units, the polyadenylation cleavage site, and transcription continuing past the polyadenylation cleavage site (**Figure 1A**). Since PRO-seq data provides strand-specific measurements of RNA polymerase, labels were separately enumerated on the plus and minus strand. Training examples for each label were selected using heuristics based on a variety of data, including GENCODE annotations, GRO-cap peak calls to indicate TIRs and TSSs, poly-adenylation site enriched RNA-seq data to identify polyadenylation cleavage sites, and CAGE which, together with GRO-cap, separated stable and unstable TSSs (**Table 1**). Input to the CNN consisted of two vectors representing the PRO-seq signal on the plus and minus strand in 50bp non-overlapping windows over a total genomic region of approximately 50 kilobases. Free parameters, including the non-overlapping window size, total genomic area provided to the CNN, and the size and depth of the CNN itself, were optimized to achieve the highest accuracy based on the area under the precision recall curve (auPRC) (see Methods).

**Figure 1:**
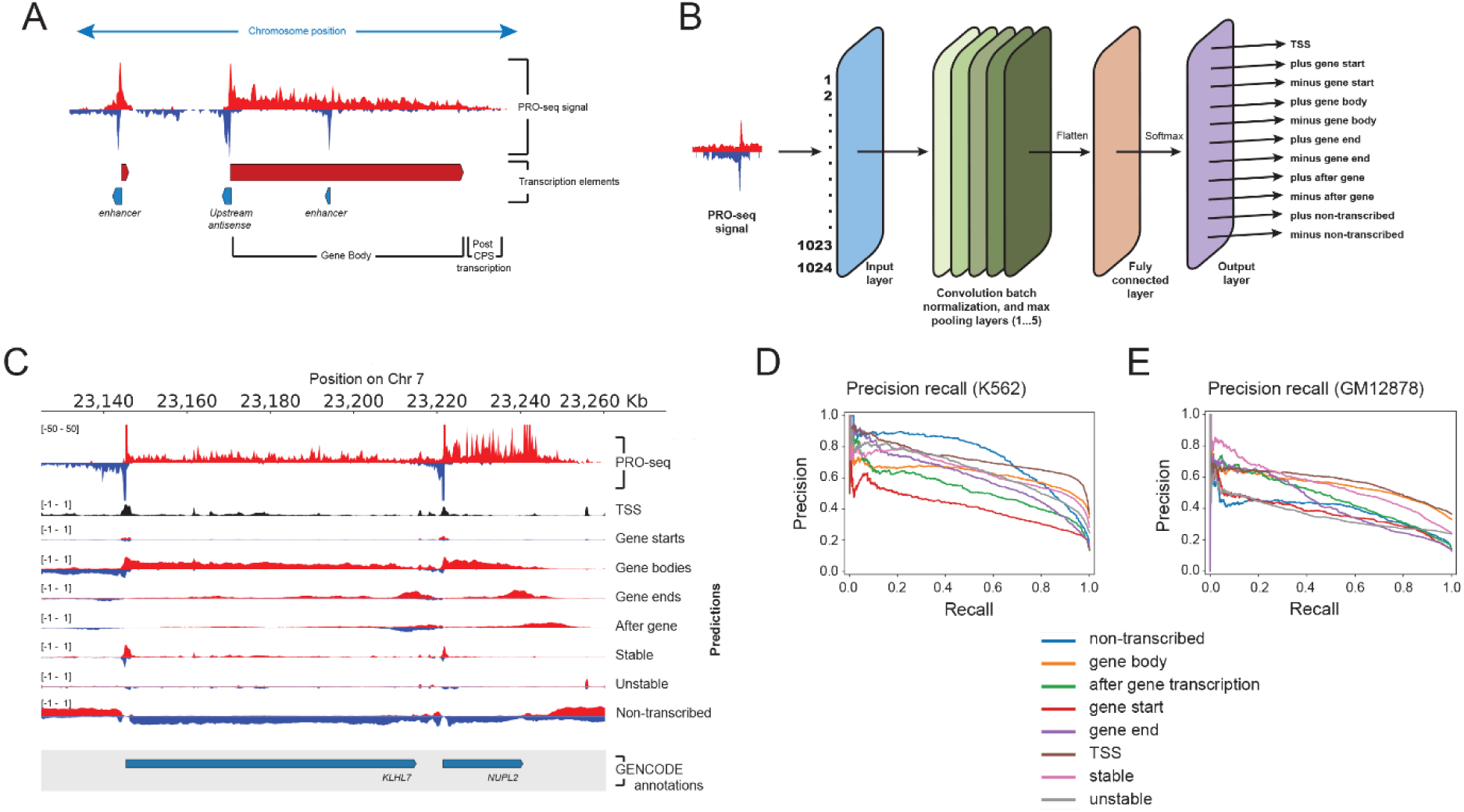
Experimental design, implementation, and performance of a convolutional neural network (CNN) designed to predict features in PRO-seq data. (A) Depiction of PRO-seq signal showing how the various shapes it produces correspond to different transcription element annotations. (B) Schematic depicting the basic structure of the CNN composed of a 1024 x 2 input layer, 5 convolutional layers, each interspersed with a batch normalization layer and a MaxPool layer, and a fully connected output layer. (C) An example PRO-seq signal aligned with the output CNN predictions at the *KLHL7* locus on chromosome 7. There is good correspondence between features in the PRO-seq signal and predicted features derived from the CNN. To assess the performance of the CNN, we employed several methods for testing the predictive capability of the CNN. For example, in (D) and (E) we determined the Precision recall curves for the CNN predictions for K562 (area under the curve values range from 0.32 to 0.64) and GM12878 respectively.

The best performing model, which we call <u>c</u>onvolutional discovery of <u>g</u>ene <u>a</u>natomy using <u>P</u>RO-seq (CGAP), contained an input layer of 2x 1024 windows, five convolutional and max pool layers, a fully connected layer, and an output layer of 15 nodes representing each of the pre-defined labels (**Figure 1B**). We trained CGAP using approximately 740,000 genomic windows and corresponding labels in K562 cells. The K562 cell line was chosen because it is an ENCODE Ter 1 line and also includes a large number of high quality PRO-seq datasets. To avoid learning features that reflect batch effects of a single PRO-seq library, we used data from four different GRO-seq and PRO-seq datasets, with two PRO-seq datasets held out as validation and test sets (see G1 through G7 in **Table 1**). Since the characteristic shape associated with each of the labels of interest is similar in different cell types, we assume that a model trained in K562 will be transferrable to any mammalian cell line in which features of gene and transcription unit structure are conserved.

We evaluated the accuracy of CGAP on held-out K562 and GM12878 datasets. Visualizing data on a genome browser showed a reasonably good correspondence between labels and the expected features in the PRO-seq signal (**Figure 1C**). Gene bodies tended to increase in signal at the transcription start site and remained consistently high until near the polyadenylation cleavage site, at which point gene body signal trailed off as the post poly(A) signal increased. Likewise, the non-transcribed label was highest in regions with little or no signal on that strand. Non-transcribed labels tended to be negatively correlated with both gene body and gene end labels, but often overlapped TIRs associated with unstable antisense or enhancer RNAs. The stable and unstable transcription start site states accurately marked the 5’ end of long transcription units, or short and unstable RNAs, respectively.

CGAP performed reasonably well genome-wide in both held-out K562 and GM12878 datasets (median auPRC for K562 = 0.635; [0.41 - 0.73]; median auPRC for GM = 0.445; [0.36 - 0.57]; median auROC for K562 = 0.865; [0.82 - 0.94]; median auROC for GM = 0.79; [0.64 - 0.84]; **Figure 1 D-E; Supplemental figures S2A, S2B**). We note that heuristics used to define the distinct types of transcription units are an imperfect ground-truth, and therefore the estimates of genome-wide auPRC likely underestimate accuracy. Some labels performed better in K562 than GM12878. The most notable example of a cell-type difference was the label for a non-transcribed region. Since this label largely reflects a lack of signal in the surrounding region, we attribute cell-type differences to the higher sequencing depth in the holdout K562 cell dataset compared with GM12878 (∼400 million in K562 vs. 100 million in GM12878). Finally, the stable and unstable labels performed well in classifying whether a particular transcription start site encoded a stable transcription unit, or an unstable transcription unit that was rapidly degraded. Thus, the multi-task CNN dissected the anatomy of an active transcription unit with reasonably high accuracy.

### 3.2 CGAP-HMM reads the boundaries of transcription units using CGAP landmarks

We hypothesized that labels learned by CGAP representing anatomical landmarks of transcription units would be useful in improving the accuracy and sensitivity of transcription unit detection. While the predictions made by CGAP for each 50 bp region along the chromatin were reliable on average, predictions that did not match the surrounding signal were reasonably common (for instance, a prediction of a non-transcribed region within a gene body, or *vice versa*). Thus, CGAP alone was not able to convert predictions of these 50 bp regions into contiguous transcription units spanning many kilobases.

We reasoned that we needed an additional step to merge each 50 bp region into contiguous regions of transcribed or non-transcribed chromatin. This additional step needed to satisfy two major criteria: First, it needed to act as a smoothing function for CGAP signal, ignoring changes in CGAP state that were either small in magnitude or not sustained across broader regions. Second, recognizing that gene annotations are at best a noisy representation of the transcription occurring in any given cell type, we needed an approach that could be fit robustly without a gold-standard training set.

We settled on using a hidden Markov model (HMM) for this task (Durbin, 1998). We compared HMMs to several discriminative methods, including a random forest classifier, an adaptive boosting classifier, a gradient boosting classifier, and a multilayer perceptron (Pedregosa *et al*., 2011), across a set of metrics used to evaluate the accuracy of transcription unit boundaries as well as manual inspection of transcription unit predictions (see Methods). Not surprisingly, the decision tree-based algorithms performed worst since these models were not developed for data smoothing. The multilayer perceptron approach also severely underperformed the HMM, likely due to the unreliable training data. We did not try applying recurrent neural networks, which are more commonly used in time series classification problems, but we predict these would have similar problems to the multilayer perceptron due to a lack of gold-standard training data. We thus adopted the HMM going forward.

The optimal HMM version modeled PRO-seq data using three separate states that represent non-transcribed regions, gene bodies, and post-poly(A) transcription (**Figure 2A**). CGAP-HMM is the combination of the predictions made by the CGAP neural network and this optimal HMM (**Figure 2B**). The labels learned by CGAP that correspond to each of these states (i.e. gene bodies) were modelled as emissions using a gamma distributed random variable. It also used the plus and minus gene start positions predicted by the CNN as co-variates to adjust the transition probability from a non-transcribed to a transcribed state (see Methods). These gene start positions are a subset of the TSS predictions, corresponding to stable TSSs. We tested many other HMM structures as well as incorporating additional CGAP labels. Several of these alternative models had interesting properties but did not perform as well in genome-wide metrics (see section *3.4 Performance comparison of CGAP-HMM, cGAN and existing transcription unit prediction tools*, below).

**Figure 2:**
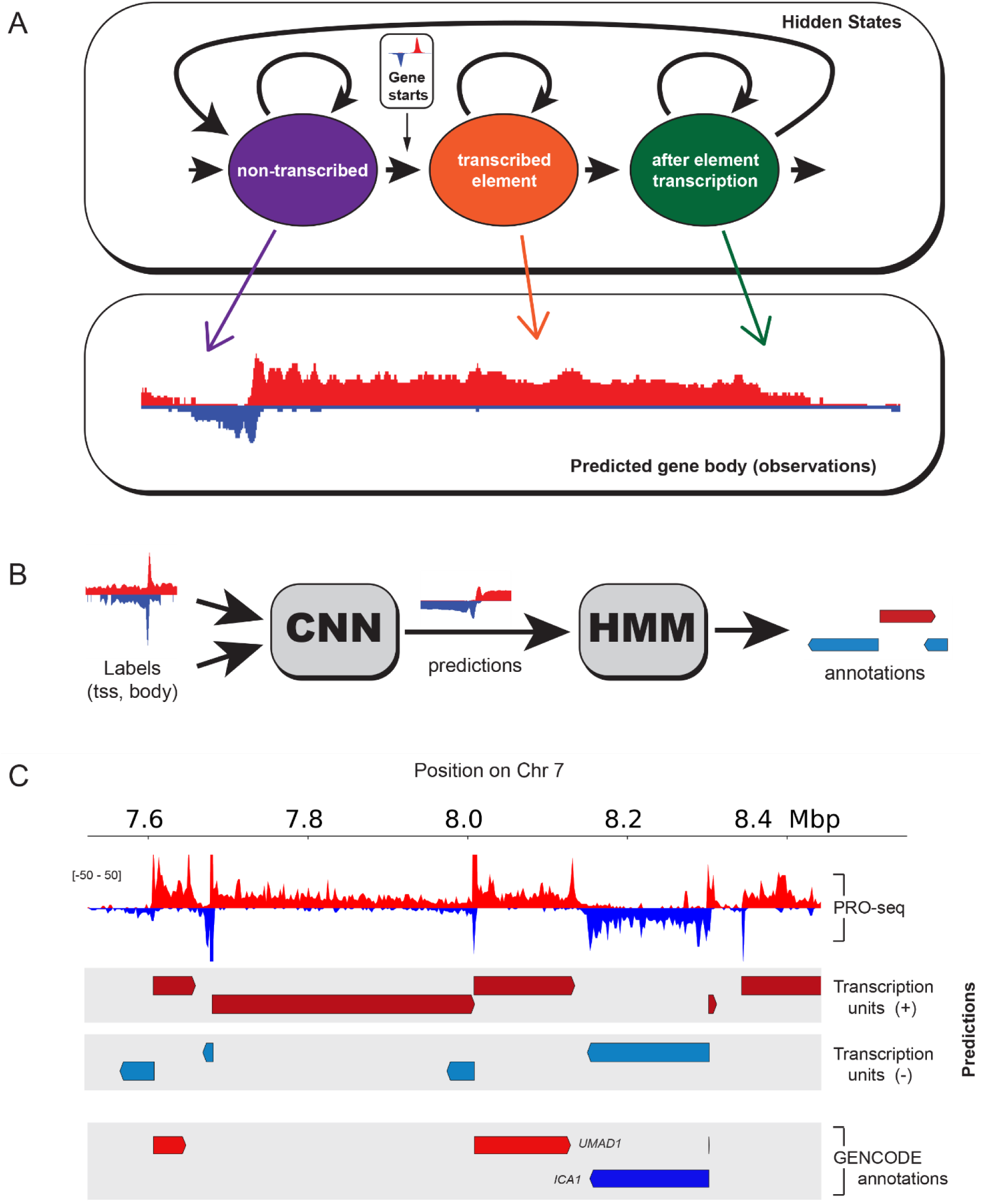
Experimental design of HMM, together with its integration with the CNN, and output predictions. (A) Schematic depicting the HMM configuration and allowed state transitions. Predictions made by the CNN for gene bodies are used as emissions. (B) Integration of the HMM with the CNN where the predictions obtained from the CNN are used as emissions and covariates for the HMM. (C) PRO-seq signal aligned with HMM predictions for the *UMAD1* locus on chromosome 7 shows good correspondence between features in the PRO-seq signal and HMM predicted annotations.

CGAP-HMM predicted boundaries of active transcription units that agreed reasonably well with the expected patterns based on both gene annotations and manual inspection. For instance, CGAP-HMM was able to separate the protein coding gene *UMAD1* from a longer transcription unit starting upstream and extending into the *UMAD1* promoter (**Figure 2C**). Likewise, CGAP-HMM recovered the location and reasonable boundaries of un-annotated transcription units in the locus surrounding *UMAD1*. Thus, to a first approximation, CGAP-HMM predicted transcription unit annotations with reasonable estimates of the boundary.

### 3.3 Predicting transcription units using cGANs

CGAP required us to specify labels of interest and provide labeled training data. While this strategy appears to work reasonably well in practice, it has several important limitations. First, labels are based on error prone heuristics obtained using molecular tools that indirectly measure the label of interest. Second, CGAP may miss additional labels that are highly informative about specific aspects of transcription unit structure, but which we did not think of *a priori*. We therefore set out to develop a generative modelling strategy which maps PRO-seq data to gene annotations, learning informative shapes using generative learning tools during the process.

We developed a conditional generative adversarial network (cGAN) which learns to map PRO-seq signal to gene annotations. A cGAN is a generalization of a generative adversarial network (GAN) (Goodfellow *et al*., 2014) which trains two separate CNNs: a generator, which generates a realistic example of data from a uniformly distributed random vector; and a discriminator, which seeks to correctly classify whether a given image was generated by the generator or is a real example of the data (see Methods). A cGAN builds on this concept by transferring specific samples from an input domain to the desired output domain. For example, cGANs introduced by (Isola *et al*., 2017) learn mappings from grayscale, outline pictures of handbags to fully colored images of the same bags, or learns mappings from satellite images to a street map view of the same area.

We used a cGAN to learn the mapping between PRO-seq signal and gene annotations. Our cGAN (**Figure 3A**) takes as input PRO-seq data in 50 bp non-overlapping windows on both the plus and minus strand over a genomic interval of 1.6 MB. We trained the cGAN to perform translations from PRO-seq signal to transcription units (**Figure 3B**). We attempted to use two sources of transcription units: first, the set of active GENCODE annotations, and second, annotations derived from an ensemble method combining the best three performing transcription unit annotation approaches (see Methods); both of which performed similarly in benchmarks. The trained cGAN takes as input PRO-seq data over a 1.6 MB genomic interval and returns a value between 0 and 1 representing the confidence that 50 bp bin is found inside of a gene annotation.

**Figure 3:**
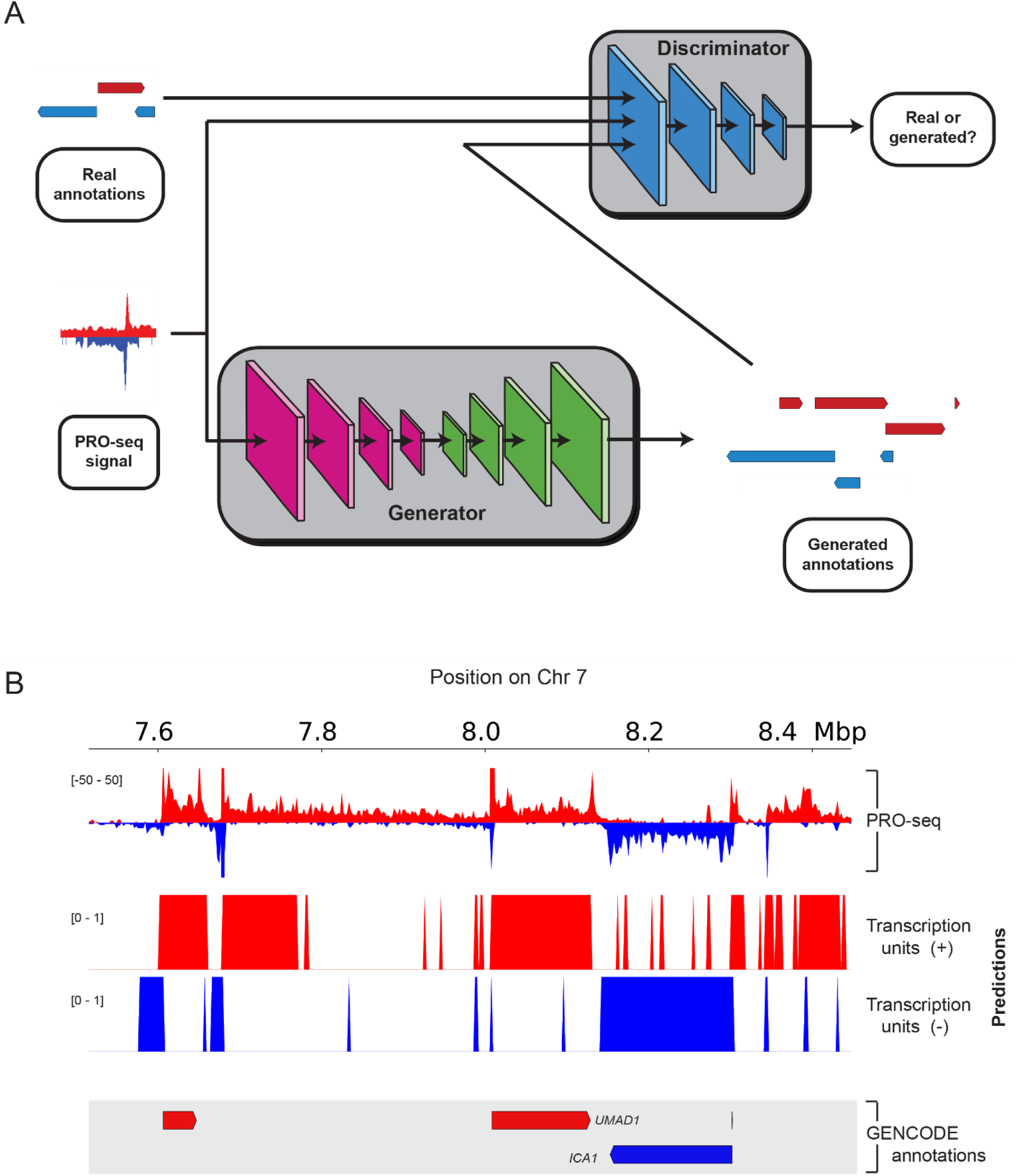
Experimental design of the cGAN and example predictions. (A) The cGAN is designed to produce generated annotations via a generator network (set up as a modified encoder-decoder). If trained correctly, the quality of the generated annotations should be at such a high level that, when fed into a discriminator (implemented as a PatchGAN with a similar set up to the encoder), the latter is unable to distinguish between the generated and the real/actual annotations. (B) Example PRO-seq signal and cGAN predictions for the *UMAD1* locus on chromosome 7 shows good correspondence with the longer, well-expressed features in the signal.

### 3.4 Performance comparison of CGAP-HMM, cGAN and existing transcription unit prediction tools

We compared CGAP-HMM and the cGAN approach to existing transcription unit identification methods by manual inspection of the UCSC genome-browser. We selected two methods (groHMM (Chae *et al*., 2015) and T-units <u>(Danko et al., 2018, GitHub:</u> https://github.com/andrelmartins/tunits/tree/master)) that use nuclear run-on assays (GRO-seq and PRO-seq respectively) to identify the boundaries of transcription units. Both T-units and groHMM use similar HMMs on raw PRO-seq to identify transcription units. groHMM was introduced using a two-state HMM, dividing the genome into transcribed and non-transcribed regions. T-units improved on groHMM by adding a third state that attempts to model transcription occurring after the end of a transcription unit, and by making the transition from the non-transcribed to the transcribed state conditional on TIRs detected using dREG. Comparison of transcription unit calls on a genome browser demonstrate the strengths and weakness of the distinct methods (**Figure 4A, Supplementary figure S4A, S4B**). T-units and groHMM tend to underestimate the lengths of transcription units with lower levels of transcription, for instance in the case of the upstream antisense transcription unit of *UMAD1*. In contrast, T-units frequently over-estimates the length of genes, incorporating post CPS transcription into the gene estimate. CGAP-HMM can miss short but highly expressed non-coding genes, such as the upstream antisense transcript associated with the non-coding transcript starting at ∼8.35MB on chr7. Nevertheless, all three transcription unit annotations performed reasonably well at most loci. In contrast, the cGAN approach tended to split each annotation into multiple parts, hugely underestimating the length of transcription units.

**Figure 4:**
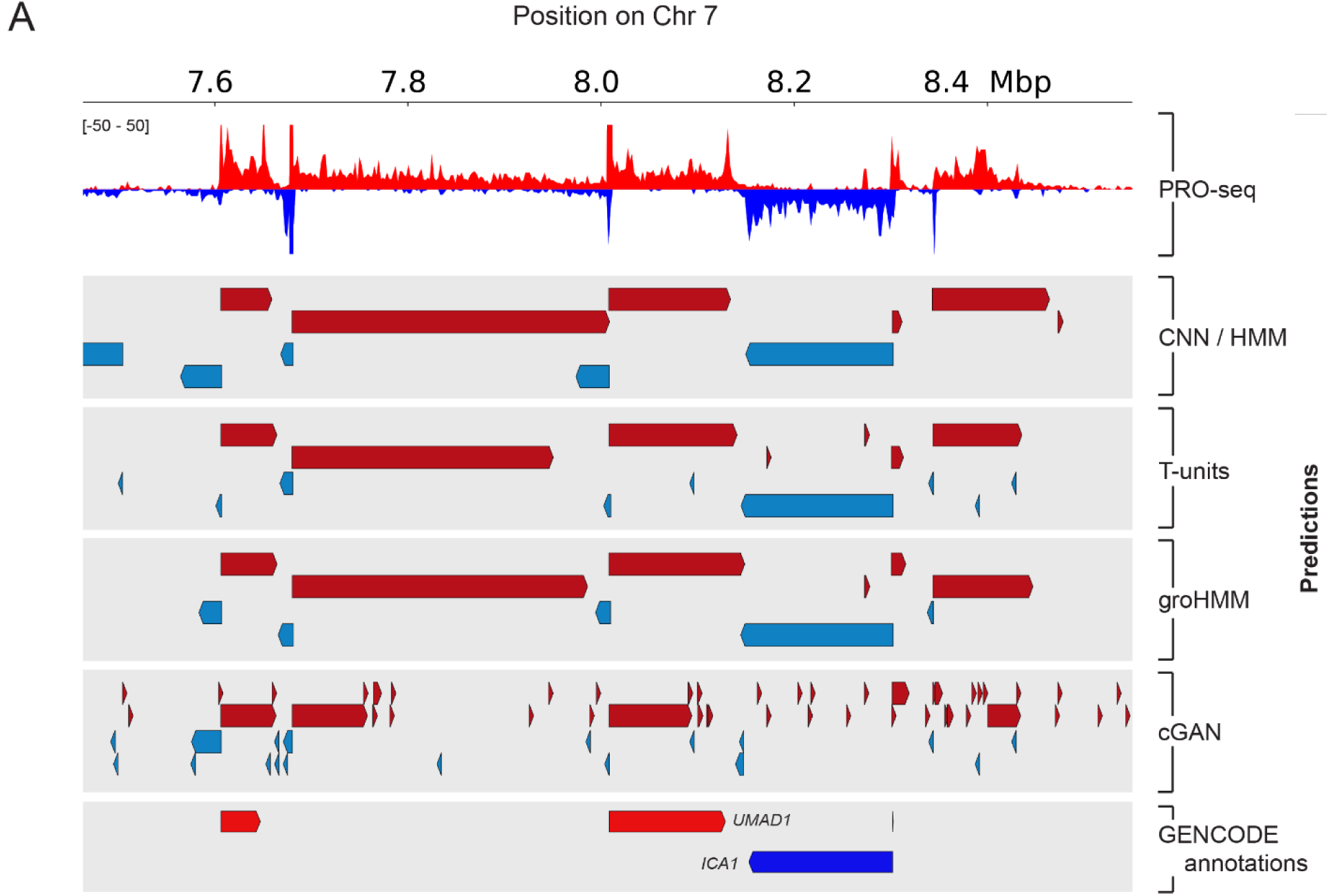
Comparison of the CNN/HMM and cGAN with other annotation methods for predicting features within PRO-seq data. T-units and groHMM predict a single gene at the *RABGEF1* locus. However, the CNN/HMM reveals that *RABGEF1* is actually comprised of many closely associated isoforms that correspond with the gaps in the CNN/HMM predictions, preceded by two distinct genes: *KCTD7* and *NR_110037*. The cGAN only partially detects these individual genes and fails to identify transcription units on the minus strand.

The differences in the performance of the CGAP-HMM, T-units, and groHMM methods provided us with the motivation to combine the three in such a way as to maintain the strengths of each while also attempting to compensate for their different weaknesses (described in section 2.7 *Building a consensus for CGAP-HMM, groHMM, and T-units* above).

We evaluated this consensus approach and all individual methods (including each of the decision tree-based and neural network approaches mentioned above in section 3.2 *CGAP-HMM reads the boundaries of transcription units using CGAP landmarks*) using several types of metrics that measure different aspects of transcription unit identification. First, we compared the rates at which each method disassociated gene annotations into separate candidate transcription units, and the rate at which separate genes were merged into a single candidate transcription unit annotation (see section *2.8 Genome-wide accuracy metrics for transcription unit identification* above; **Supplementary figures S1A, S1B**). We note that while the optimal method would get both values as low as possible, incorrect parameter tuning can result in one value decreasing at the expense of the other. For example, it would be easy to decrease the count of annotations disassociated by increasing the number of annotations merged (or vice versa), but this does not provide a good solution to the transcription unit identification problem. CGAP-HMM performed the best on this metric, splitting nearly the same low number of annotated genes as groHMM, while merging only ∼half the number. The cGAN did not perform well in this metric, merging fewer transcripts than groHMM, but disassociating significantly many more (**Figure 5A**). It should be noted that the decision tree-based and neural network approaches appeared to perform exceptionally well on this metric, but this result is misleading since these methods predicted so few annotations that the number that were merged and / or disassociated is artificially low.

**Figure 5:**
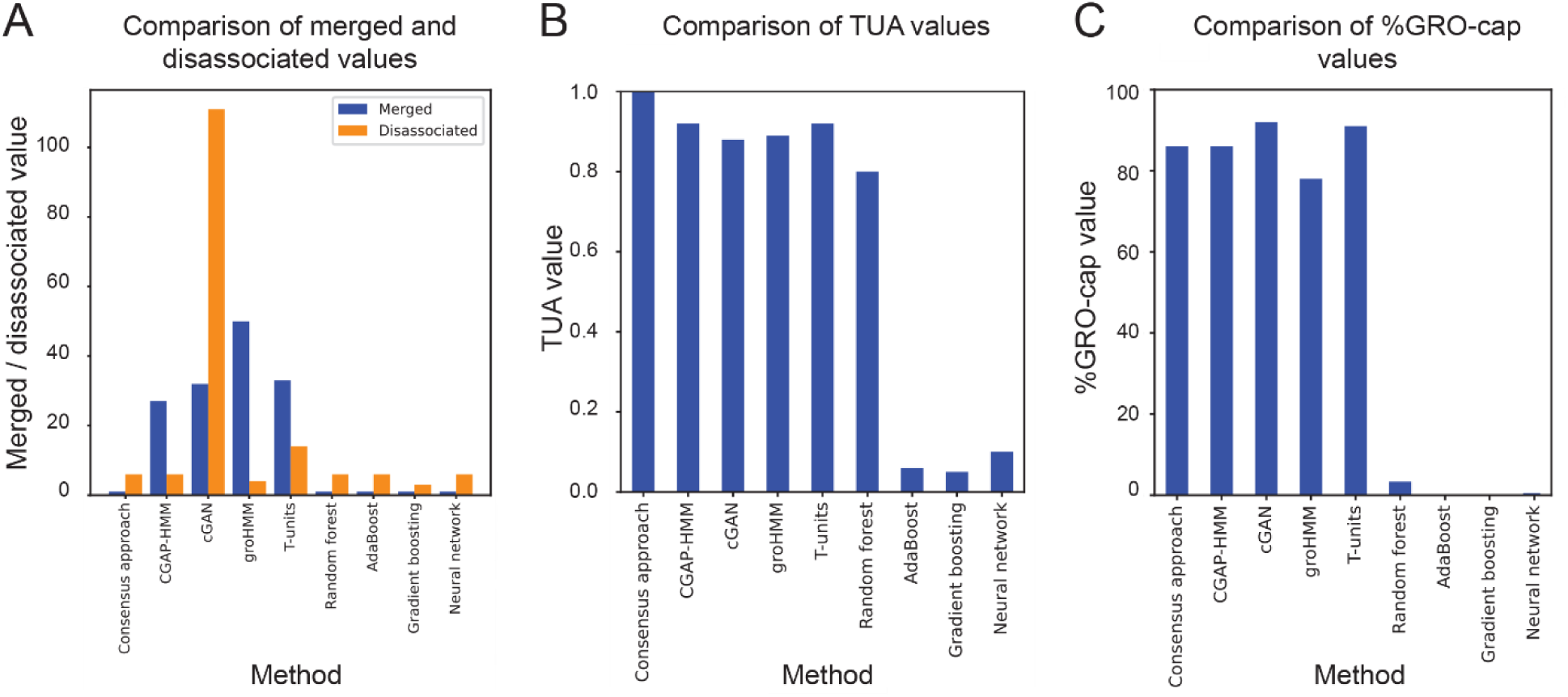
Metrics used to evaluate the performance of each of the methods. (A) Comparison of merged and disassociated values. (B) Comparison of TUA values. (C) Comparison of %GRO-cap values.

Next we examined the TUA metric (Chae *et al*., 2015), which measures the accuracy of the 5’ and 3’ end of transcription unit predictions (**Figure 5B**). The TUA metric ranges between 0 and 1, and values near 1 indicate higher agreement with the ground truth. A comparison of TUA showed that CGAP-HMM and T-units both achieved the best results, a TUA of 0.92, higher than groHMM. The cGAN did not perform well in this metric, although it did outperform the decision tree-based and neural network approaches (**Figure 5B**).

As a metric for how different TU callers handled unstable, intergenic transcripts, we examined the fraction of intergenic GRO-cap sites that were captured by each transcription unit prediction method. cGAN performed with the highest sensitivity in this task, discovering 92% of GRO-cap sites, and T-units was close behind. CGAP-HMM performed well with 86%, and groHMM also performed well at 78%.

These results show that all transcription unit prediction tools had a reasonably high sensitivity for intergenic transcription, with the exception of the decision tree-based and neural network approaches (**Figure 5C**).

Collectively, although all transcription unit identification tools performed reasonably well (aside from the decision tree-based and neural network approaches), CGAP-HMM appears to make improvements, which are in some cases substantial, on stable protein-coding genes at the expense of calling fewer very short, unstable transcription units.

The Merged, Disassociated, TUA, and % GRO-cap metrics were also used to evaluate the performance of various HMM architectures (**Supplemental Figure S5A, S5B, S5C**), ultimately leading us to select the HMM described in figure 2A; i.e. three states, (non-transcribed, transcribed, after element transcription), with a single covariate, (the gene starts predicted by CGAP), used to modify the transition probability from non-transcribed to transcribed states. The HMMs described in supplemental figure 5 are by no means an exhaustive list of the architectures we experimented with, but rather serve as examples of some of the architectures we considered. As a general rule, we found the simpler models outperformed the more complex models – while it is true that our three-state model with a single covariate did better than a two-state model, increasing the number of states beyond three and the covariates beyond one resulted in poorer performance.

### 3.5 Assessing our model on other cell types / organisms

Next, we asked whether CGAP-HMM can identify the features we have discovered in the PRO-seq signal of K562 cells in other cell types / tissue types. To this end, we ran both our pre-trained CGAP-HMM and ensemble models on five additional cell lines: GM12878, MCF-7, HeLa, HTC116, and CD4+ T-cells (**Figure 6A**). To compare the results of our analysis between these cell types we ran the same evaluation metrics as used for K562 cells (**Figure 7A, B, C**). These comparisons revealed that that method performed similarly on all cell types tested, with respect to the TUA scores. Even for a cell line that is sequenced at a much lower depth, namely the CD4 cell line, we could detect the same features using CGAP-HMM. However, values for disassociated annotations were consistently higher. We note that data for GRO-cap sites are only available for K562 and GM12878 cells, so we were unable to make the % GRO-cap recovered measurement for the remaining cell types.

**Figure 6:**
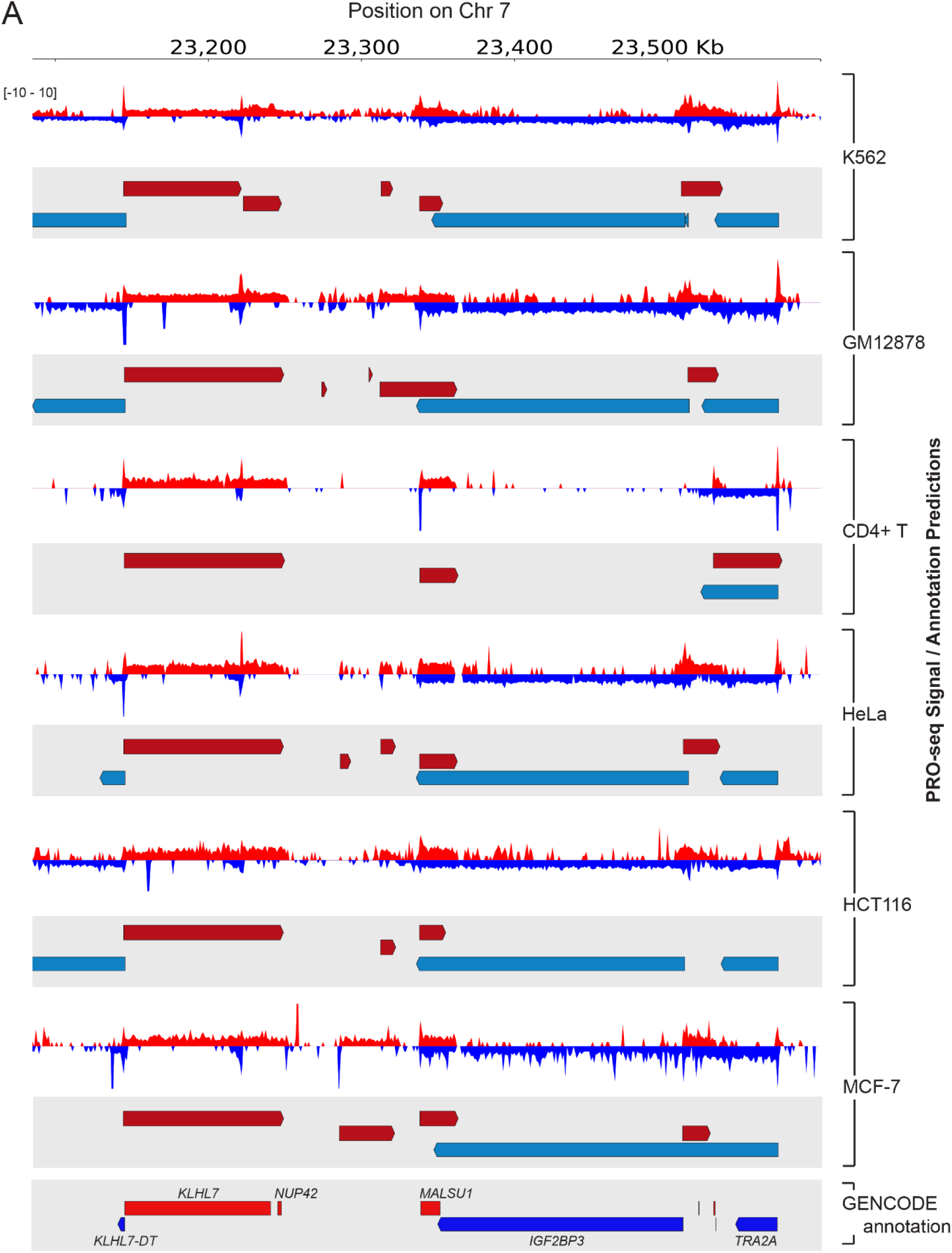
Comparison of CNN/HMM output from PRO-seq data acquired from a range of different cell types. To evaluate the overall performance of the CNN/HMM, we tested it with PRO-seq data from 5 additional cell types. Shown here are PRO-seq signals from each cell type aligned with their associated predictions for the KLHL7 locus on chromosome 7.

**Figure 7:**
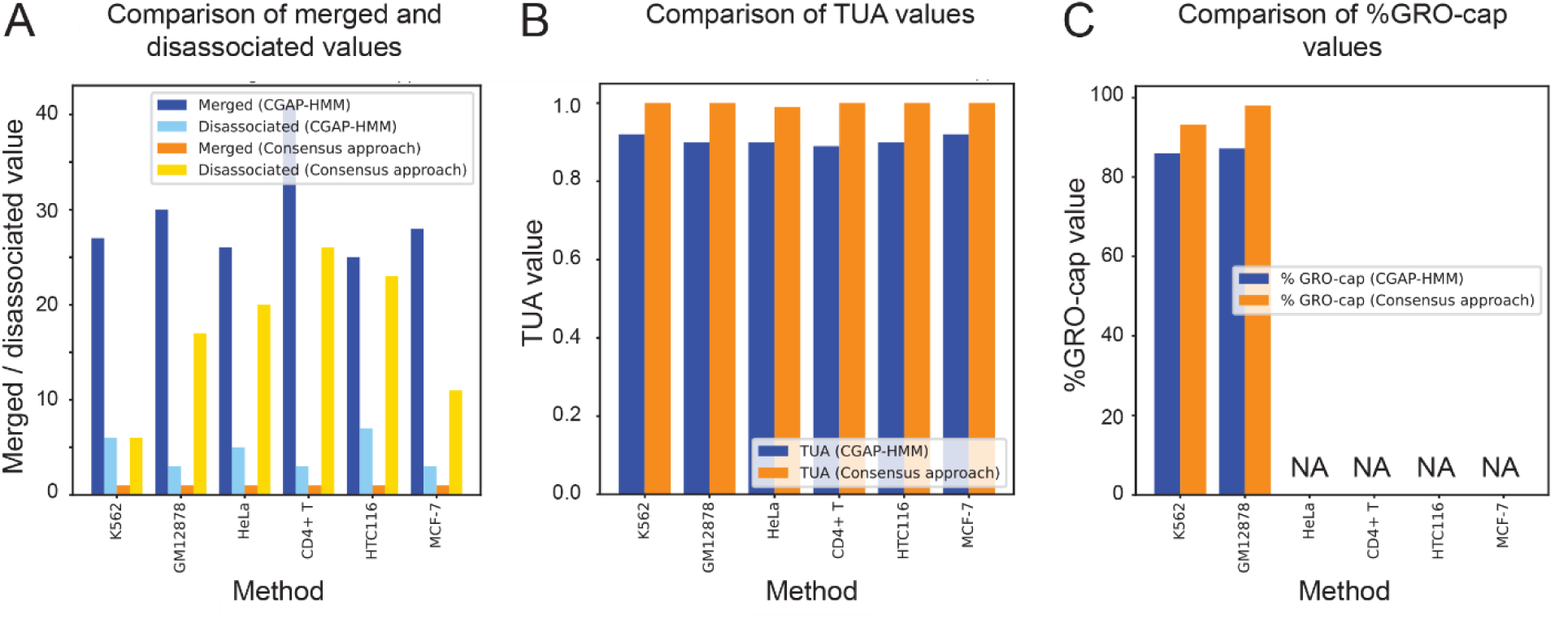
Metrics used to evaluate the performance of CGAP-HMM vs. the consensus approach for each cell type. (A) Comparison of merged and disassociated values. (B) Comparison of TUA values. (C) Comparison of %GRO-cap values.

## 4. DISCUSSION

Gene annotators have been built by applying HMMs directly to PRO-seq data without the intermediate step of feature detection / pattern recognition provided by a neural network. We show that by combining the feature detection strengths of neural networks with HMMs we can get improved annotations.

We used two novel approaches, first a CNN coupled with an HMM, and second a cGAN. The neural network approach was chosen because these are one of several machine learning techniques that are ideal for handling the non-linearities inherent in the task of translation from PRO-seq data to genome annotations. Indeed, we demonstrate here that these methods can detect subtle patterns of RNA pol II activity. For example, we were able to model the after-gene transcription that occurs when RNA pol II continues to elongate beyond the 3’ end of transcriptional units.

When we conceived the cGAN approach, it was with the aim of producing the distinct begin and end boundaries necessary to call annotations, without the need for additional smoothing by an HMM. In this regard, the cGAN was partially successful in that it was able to identify the start of transcription units, but it did not do as well as other methods in detecting the end points, failing to distinguish transcription after unit boundaries from transcription with the body of a unit body. It also performed poorly, in most cases, in picking up shorter transcripts (such as putative enhancers), and consequently ranks at the bottom of the comparisons we made with other methods.

For the CNN, we adopted the approach of producing continuous distributions for each of the features we were attempting to detect, and then employed an HMM to determine the most likely set of annotations given those distributions. Not surprisingly, considering the transcriptional noise inherent in the PRO-seq signal and consequent noisiness of the CNN’s predictions, this additional smoothing step gave us better results.

Overall, our data indicate that both the CNN coupled with an HMM, and the cGAN have great potential. For the CNN / HMM approach the simple addition of more training data with better labels would improve performance. It should also be noted that supplying the HMM with data from additional assays (rather than relying on the CNN’s predictions for start and end sites) improves its performance (data not shown). Our original intent was to build a method that could work in organisms where assay data is limited, but if this data is available then improved annotations can be obtained by including it. In the case of the cGAN, many of the problems we encountered in training are still active areas of research, so this approach may depend on future developments in GAN technology before it becomes a viable method for transcription unit annotation.

## 5. DATA ACCESS

## ACKNOWLEDGMENTS

We thank André Martins and members of the Danko and Siepel labs for thoughtful discussions and suggestions about this work. CGD, PRM, and JC acknowledge support provided by an NHGRI (National Human Genome Research Institute) R01-HG009309 grant to CGD. The content is solely the responsibility of the authors and does not necessarily represent the official views of the NHGRI.

## Supplemental figures

**Supplemental Figure 1:**
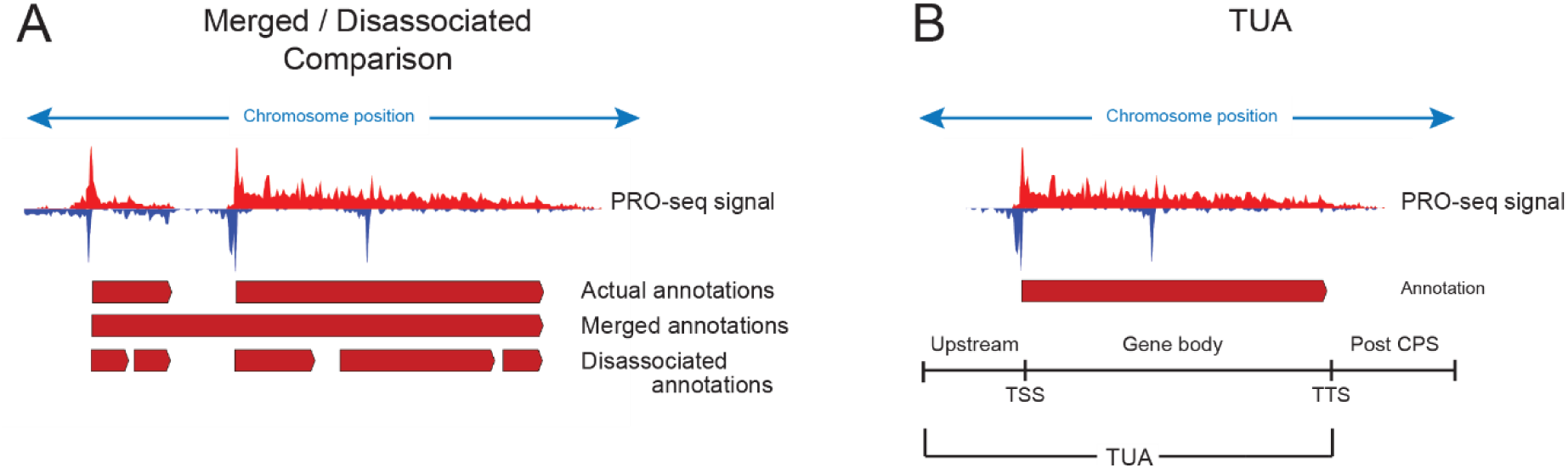
Depiction of metrics used in the evaluation of transcription unit (TU) boundaries. (A) Merged / disassociated TU boundaries. Different methods can erroneously break annotated TUs into smaller predicted fragments, or alternatively a single TU prediction can erroneously merge two or more separate TU annotations. (B) TUA metric. This metric quantifies TU annotation error in either over-or under-estimating the length of gene annotations. It measures the accuracy of the 5’ and 3’ end of transcription unit predictions by dividing each prediction into three sections (upstream of the TSS, within the transcribed region, and downstream of the polyadenylation cleavage site). The TUA metric will penalize predicted annotations that begin before the TSS of the ground truth annotation, or that end before the polyadenylation cleavage site. It does not penalize for extending annotations past the polyadenylation cleavage site. The TUA metric ranges between 0 and 1, and values near 1 indicate higher agreement with the ground truth.

**Supplementary Figure 2:**
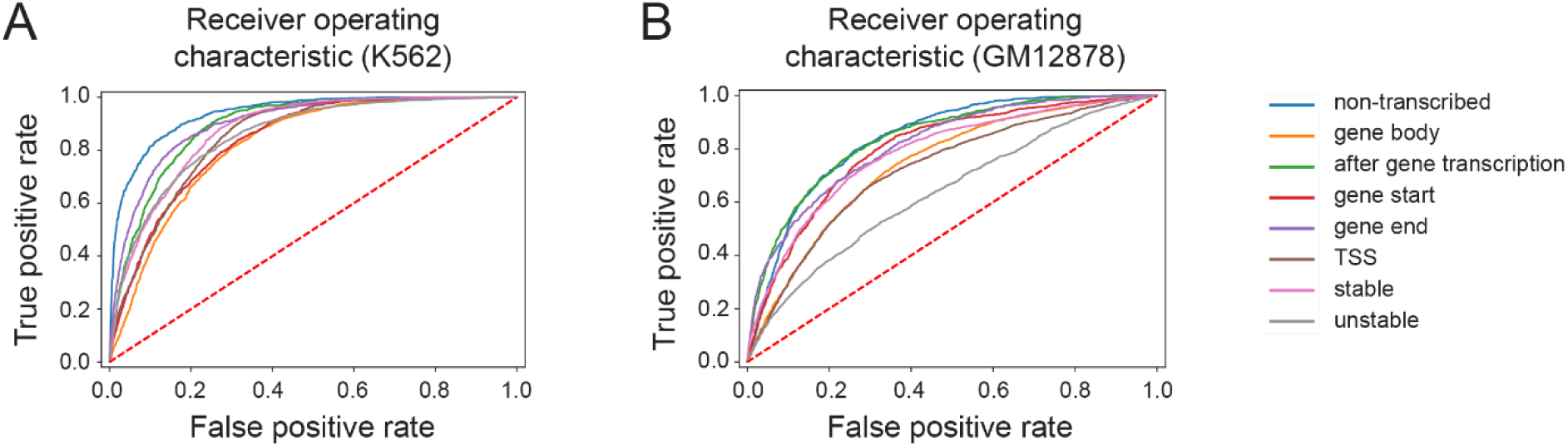
Receiver Operating Characteristic curves for CNN predictions. (A) Results for K562 (area under the curve values range from 0.83 to 0.95). (B) Results for GM12878.

**Supplemental figure 4:**
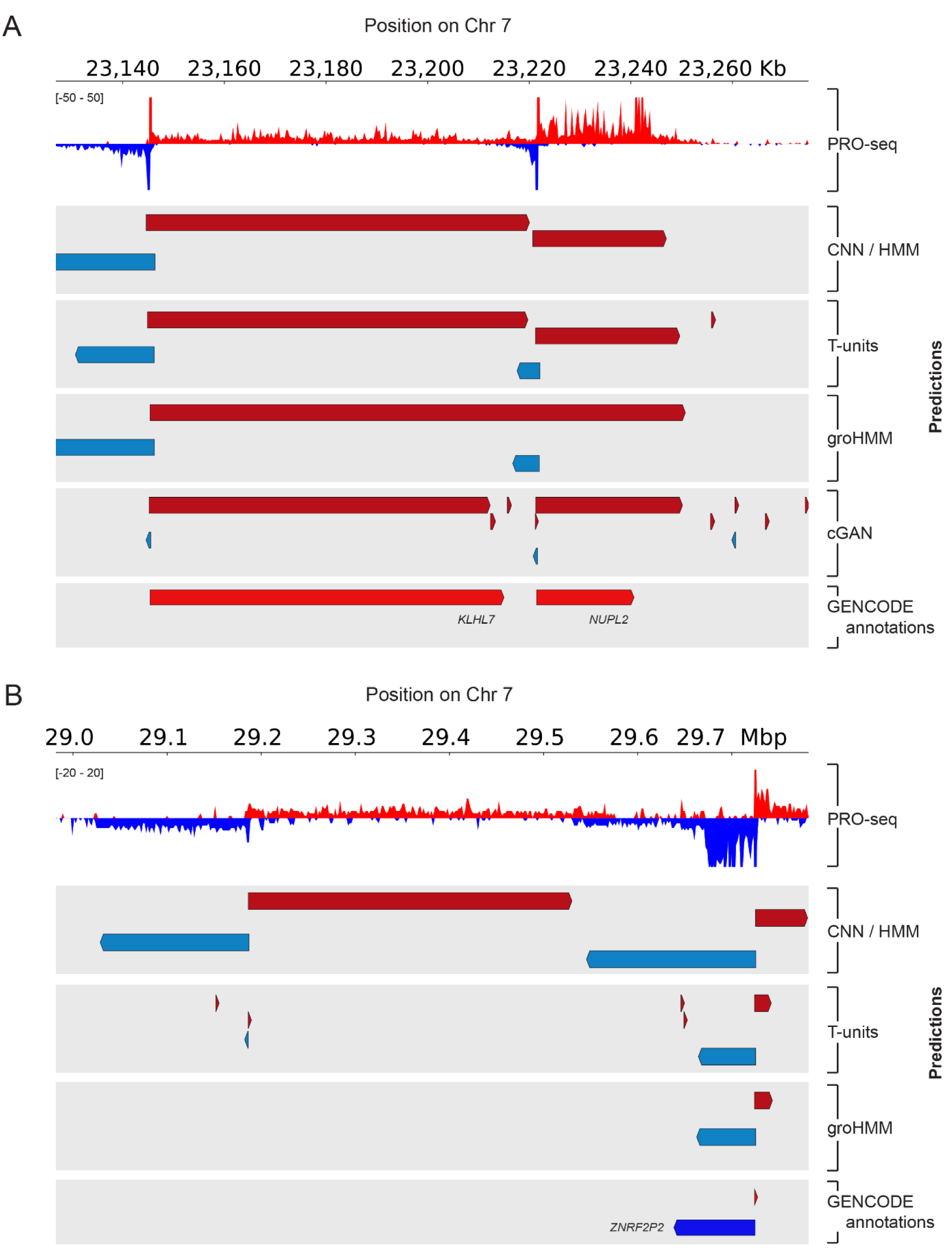
Comparison of the CNN/HMM and cGAN with other annotation methods for predicting features within PRO-seq data. (A) All methods show reasonably good correspondence with the GENCODE annotations at the KLHL7 locus. (B) T-units and groHMM predict a single transcription unit at the *ZNRF2P2* locus. However, the CNN/HMM reveals other possible transcription units that can be detected by visual inspection of the PRO-seq signal.

**Supplementary figure 5:**
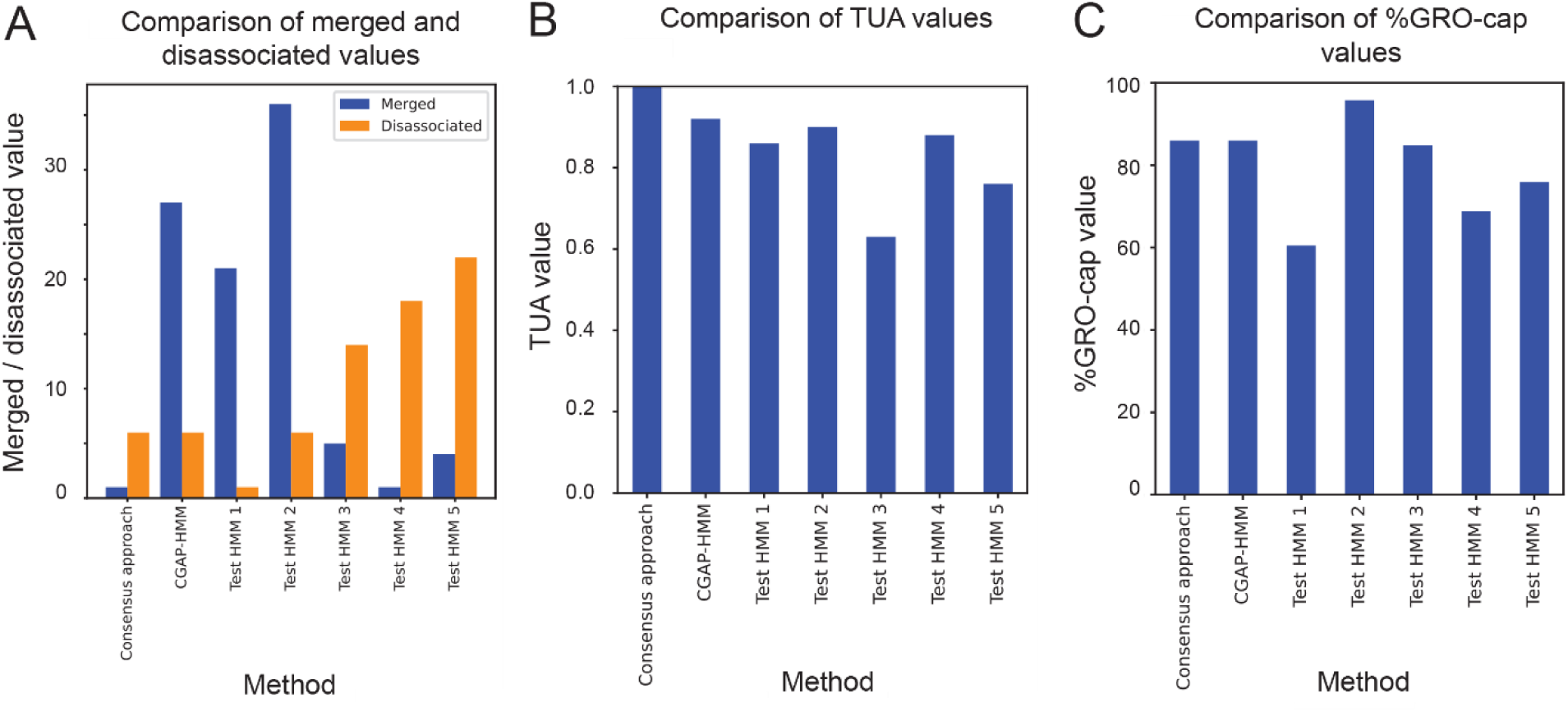
Metrics used to evaluate the performance of various HMM architectures. (A) Comparison of merged and disassociated values. (B) Comparison of TUA values. (C) Comparison of %GRO-cap values.

